# Chromatin Condensation Fluctuations Rather than Steady-State Predict Chromatin Accessibility

**DOI:** 10.1101/365700

**Authors:** Nicolas Audugé, Sergi Padilla-Parra, Marc Tramier, Nicolas Borghi, Maïté Coppey-Moisan

## Abstract

Chromatin accessibility to protein factors is critical for genome activities. Dynamic changes in nucleosomal DNA compaction and higher order chromatin structures are expected to allow specific sites to be accessible to regulatory factors and the transcriptional machinery. However, the dynamic properties of chromatin that regulate its accessibility are poorly understood. Here, we took advantage of the microenvironment sensitivity of the fluorescence lifetime of EGFP-H4 histone incorporated in chromatin to map in the nucleus of live cells the dynamics of chromatin condensation and its direct interaction with a tail acetylation recognition domain (the double bromodomain module of human TAF_II_250, dBD). We reveal chromatin condensation fluctuations supported by mechanisms fundamentally distinct from that of condensation. Fluctuations are spontaneous, yet their amplitudes are affected by their sub-nuclear localization and by distinct and competing mechanisms dependent on histone acetylation, ATP, and both. Moreover, we show that accessibility of acetylated histone H4 to dBD is not restricted by chromatin condensation nor predicted by acetylation, rather, it is predicted by chromatin condensation fluctuations.

**Significance:** In higher eukaryotes, the structure and compaction of chromatin are considered as barriers to genome activities. Epigenetic marks such as post-translational modifications of histones can modify the structure and compaction of chromatin. The accessibility of protein factors to these epigenetic marks is therefore of paramount importance for genome activities. We reveal chromatin condensation fluctuations supported by mechanisms fundamentally distinct from that of condensation itself. We show that accessibility of acetylated histone H4 to double bromodomains is not restricted by chromatin condensation nor predicted by acetylation, rather, it is predicted by chromatin condensation fluctuations.

**Classification:** Biological Sciences, Cell Biology

## INTRODUCTION

The regulation of multi-level chromatin compaction required for transcriptional activity in eukaryotic cells involves post-translational modifications of histones and their specific interactants, as well as histone chaperones and ATP-dependent chromatin remodelers. Histone acetylation was the first type of such modifications to be associated with gene transcription (1). Acetylation of histone, likely through the neutralization of positive charges, decreases its affinity to DNA (2), alters nucleosome conformation (3), promotes the association of transcription factors with nucleosomes (4), and causes chromatin decondensation (5, 6), thereby providing possible mechanisms contributing to transcriptional activity. New probes now allow real-time imaging of histone acetylation (7). Yet, histone acetylation predicts the potential for gene transcription, rather than transcription itself.

Hypersensitivity to nucleases (8) has been the historical proxy to assess nucleosome organization and chromatin accessibility, but this, and derived approaches do not inform on chromatin dynamics in live cells (9). Advances in fluorescence microscopy techniques, however, have allowed to probe the intranuclear milieu permeability to presumably inert fluorescent particles and the mobility of chromatin interactants (10–14), chromatin motions (15–19), or chromatin-bound DNA torsional dynamics (20). These studies have revealed the existence of multi-scale chromatin dynamics, which relations to chromatin acetylation and accessibility remain poorly understood.

Here, we sought to directly assess the relationship between chromatin dynamics and accessibility, and its dependence on histone acetylation and ATP-dependent mechanisms. To do so, we leveraged in live cells the sensitivity to the local environment of a fluorophore fluorescence lifetime, which was previously shown to report chromatin condensation states in fixed cells (15). We combined this with direct assessment by FRET (Förster resonance energy transfer) of the interaction between acetylated H4 histone and human TAF_II_250 double bromodomain (21, 22), a module conserved in histone acetyltransferases, bromodomain and extra-terminal domain protein families of transcriptional regulators, and ATP-dependent chromatin remodeling factors (23, 24).

A detailed spatiotemporal analysis under hyper-acetylation, ATP depletion, and the combination of both conditions revealed spontaneous chromatin condensation fluctuations that are affected by distinct and competing mechanisms dependent on acetylation, ATP, and both, and by their sub-nuclear localization. Moreover, the amplitude of these fluctuations predicts chromatin accessibility, which is not restricted by chromatin condensation.

## RESULTS

### EGFP-H4 fluorescence lifetime reports chromatin condensation and its global and local heterogeneities

We imaged EGFP-H4 in live HEK cells. 80 % of EGFP-H4 was immobile as assessed by FRAP (Fluorescence recovery after photobleaching) (Fig. S1A), supporting it was mainly incorporated in chromatin. The time-averaged fluorescence lifetime *τ*̃ (see SI Materials and Methods) was slightly but significantly lower for EGFP-H4 than for EGFP alone, showing that tagged EGFP was sensitive to the presence of histones (Fig. 1A, B).

**Figure 1:**
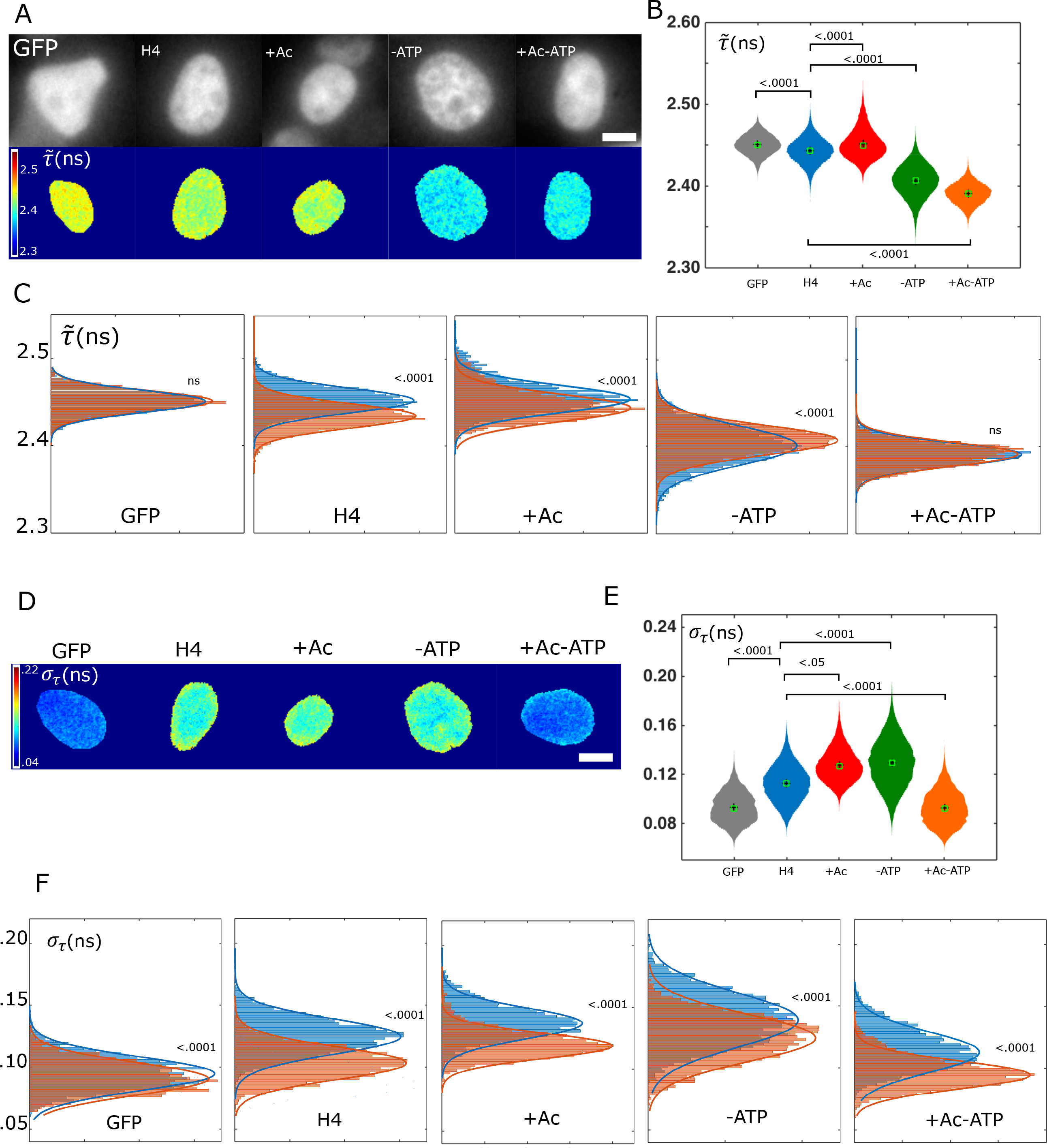
**A:** Typical images of HEK cells expressing either free EGFP or EGFP-H4 in untreated (H4), hyper-acetylation (+Ac), ATP-depletion (-ATP), or both (+Ac-ATP) conditions. Top: steady-state fluorescence intensity, bottom: time-averaged fluorescence lifetime *τ*̃. Bar = 5 μm. **B:** Time-averaged fluorescence lifetime *τ*̃ distributions in conditions above: EGFP (12 694 pixels, 7 cells); EGFP-H4 (29 288 pixels, 15 cells); +Ac (17 405 pixels, 9 cells); -ATP (32 425 pixels, 13 cells); +Ac-ATP (20 795 pixels, 8 cells). Black cross: mean, green box: median. Kruskal-Wallis test. **C:** Time-averaged fluorescence lifetime *τ*̃ distributions in conditions above for peripheral and central pixels and their Gaussian fits. Extra sum-of-squares F test with same fit for both distributions as null hypothesis. ns=not significant. **D:** Typical images of fluorescence lifetime standard deviation *σ*_*τ*_ in HEK cells expressing either free EGFP or EGFP-H4 in untreated (H4), hyper-acetylation (+Ac), ATP-depletion (-ATP), or both (+Ac-ATP) conditions. Bar = 5 μm. **E:** Distributions of fluorescence lifetime standard deviation *σ*_*τ*_ in conditions above, same pixel and cell numbers as in Fig. 1. Black cross: mean, green box: median. Kruskal-Wallis test. **F:** Distributions of fluorescence lifetime standard deviation *σ*_*τ*_ in conditions above for peripheral and central pixels and their Gaussian fits. Extra sum-of-squares F test.

To validate that *τ*̃ reports chromatin condensation, we exposed cells to the histone deacetylase inhibitor sodium butyrate for histone hyper-acetylation (+Ac)(Fig. S1B), or sodium azide and deoxy-glucose for ATP depletion (-ATP), which induce chromatin decondensation or condensation, respectively (5, 15). In +Ac condition, *τ*̃ of EGFP-H4 appeared mildly but significantly higher than in untreated condition. In –ATP condition, *τ*̃ was substantially and significantly lower than in untreated condition (Fig. 1B). These results confirm that *τ*̃ decreases with chromatin condensation. Remarkably, ATP depletion reversed the effect of hyper-actetylation: the +Ac-ATP condition exhibited a *τ*̃ even lower than the -ATP condition only (Fig. 1B). This reveals a functional interaction between histone acetylation state and ATP-dependent mechanisms.

Since *τ*̃ was broadly distributed in most conditions, we next examined whether these distributions contained spatially distinct sub-populations of condensed chromatin. We defined peripheral and central nuclear regions (Fig. S1C, see SI Materials and Methods), both of which exhibited normal distributions of *τ*̃ across pixels that were indistinguishable from each other for free EGFP (Fig. 1C). In contrast, *τ*̃ of EGFP-H4 was on average significantly higher at the nucleus periphery than in the center (Δ*τ*̃ ~ 0.02ns). Note that *τ*̃ was not normalized between cells, which implies that *τ*̃ intercellular variability is negligible in comparison to intranuclear variability. In +Ac conditions, *τ*̃ was also higher at the periphery than in the center (Δ*τ*̃ > 0.01ns). In ATP-depleted cells in contrast, differences in *τ*̃ between peripheral and central regions were abolished for hyper-acetylated chromatin (+Ac-ATP), or even reversed, *τ*̃ higher in the center than at the periphery (Δ*τ*̃ ~ - 0.007ns) in the absence of hyper-acetylation (-ATP). Altogether, these results show that chromatin is actively less condensed at the nucleus periphery.

To further characterize chromatin condensation spatial heterogeneities independently of the radial localization, we measured the spatial autocorrelation of the instantaneous lifetime *τ* (see SI Materials and Methods). For EGFP-H4, *τ* exhibited an autocorrelation length *l* of 300nm, significantly higher than that of GFP alone (250nm, the spatial resolution limit) (Fig. S1D), supporting the existence of ~600nm-diameter domains of similar condensation state. Hyper-acetylation slightly but significantly increased the size of these domains compared to that in untreated cells. In contrast, ATP depletion had no significant effect, unless in hyper-acetylated condition, where it significantly decreased the size of condensation domains below the resolution limit. The opposite effects of hyper-acetylation as a function of ATP levels on *l*, consistent with that on *τ*̃ further supports that histone state and cell metabolism interact to control condensation domain size: hyper-acetylation favors large domains of similar condensation in an ATP-dependent fashion, while it hinders the size of such domains in the absence of ATP.

### Fluctuations of chromatin condensation depend on sub-nuclear localization, histone acetylation and metabolic state

Since chromatin condensation exhibited spatial heterogeneities, we next examined whether and how it fluctuated in time. To quantify this, we measured the standard deviation of EGFPH4 fluorescence lifetime temporal fluctuations, *σ*_*τ*_ (see SI Materials and Methods). In untreated condition, *σ*_*τ*_ of EGFP-H4 was significantly higher than that of GFP alone, supporting that these fluctuations were specific to the environment of histones (Fig. 1D,E). Remarkably, ATP depletion increased *σ*_*τ*_ compared to untreated cells, which indicates that fluctuations are spontaneous and hindered by ATP-dependent mechanisms. Hyper-acetylation in ATP-depleted condition (+Ac-ATP) decreased *σ*_*τ*_ compared to ATP-depletion alone (-ATP), showing that ATP-independent, hyper-acetylation-dependent mechanisms also hinder spontaneous fluctuations. However, addition of both ATP and hyper-acetylation (+Ac condition) had on spontaneous fluctuations (-ATP condition) the mildest decrease compared to addition of any of the two (H4 untreated or +Ac-ATP). This supports the existence of ATP- and hyper-acetylation-dependent mechanisms that out-compete the effects on spontaneous condensation fluctuations of mechanisms that only depend on ATP or hyper-acteylation.

We next examined the spatial organization of chromatin condensation fluctuations. For EGFP alone, *σ*_*τ*_ was slightly higher at the periphery than in the center of the nucleus (Δ*σ*_*τ*_ < 0.005ns), showing that fluorescence lifetime fluctuations were somewhat sensitive to nuclear localization regardless of protein tagging. Nevertheless, *σ*_*τ*_ of EGFP-H4 in all conditions exhibited between nuclear periphery and center a significant difference that was at least twice larger than that of EGFP alone (Δ*σ*_*τ*_ > 0.01ns) (Fig. 1F), consistent with chromatin condensation fluctuations at the periphery larger than in the center of the nucleus, in all conditions. Thus, chromatin condensation and its fluctuations both exhibit radial organizations within the nucleus. Nevertheless, the radial organizations of condensation and its fluctuations are distinctly regulated since they respond differently upon acetylation and metabolic perturbations.

### Cell metabolism is required for chromatin de-condensation to correlate with its fluctuations

To clarify the relation between chromatin condensation and its fluctuations, we directly assessed their level of correlation in a pixel-by-pixel, non-spatially biased approach (Fig. 2A). While EGFP did not exhibit a correlation between *τ*̃ and *σ*_*τ*_ (r= - 0.1759), EGFP-H4 did in untreated condition (r=0.5318) as well as in +Ac condition (r=0.4829). Compared to untreated cells, ATP depletion led to loss of correlation (r=0.0942) within a hyper-acetylated background, and to weak anti-correlation without (r= - 0.2447). Thus, correlation between *τ*̃ and *σ*_*τ*_ is largely acetylation-independent but strongly ATP-dependent. Correlation reversal upon ATP depletion further confirms that ATP-dependent mechanisms promoting chromatin decondensation simultaneously hinder spontaneous chromatin fluctuations.

**Figure 2:**
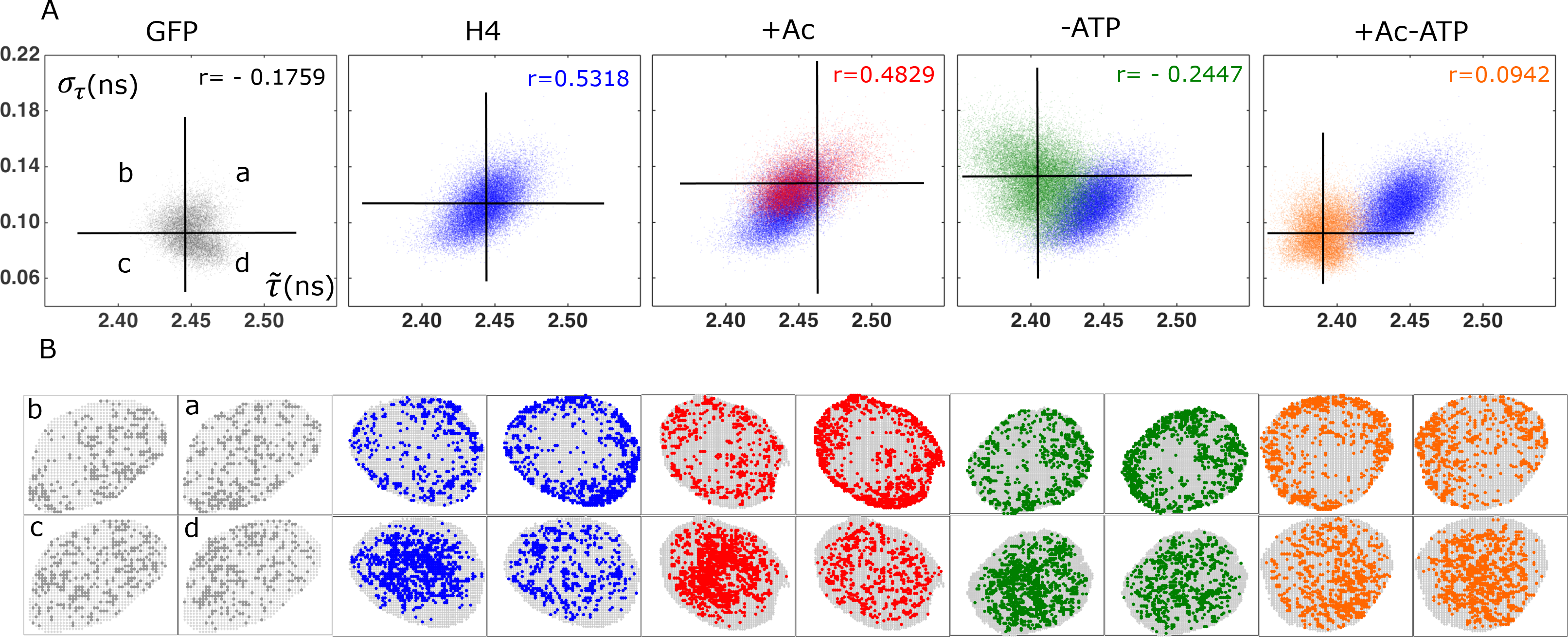
**A:***σ*_*τ*_ − *τ*̃ plots for free EGFP or EGFP-H4 in untreated (H4), hyper-acetylation (+Ac), ATP-depletion (-ATP), or both (+Ac-ATP) conditions and their respective correlation coefficients. Same pixel and cell numbers as in Fig. 1. Values for untreated cells are displayed in blue along all other conditions for visual comparison. Horizontal and vertical lines cross at *σ*_*τ*_ and *τ*̃ means. **B:** Typical localization in a nucleus of pixels from the four quadrants defined above.

To assess how this correlation was related to chromatin localization, we split the pixel population in four quadrants around the mean *τ*̃ and *σ*_*τ*_ and mapped them on the nucleus (Fig. 2B). For EGFP alone, pixels of all quadrants distributed throughout the nucleus in both peripheral and central regions. For EGFP-H4 in contrast, pixels from high *σ*_*τ*_ quadrants (a, b) were largely excluded from the central region, and pixels from the low *τ*̃ and *σ*_*τ*_ quadrant (c) were more abundant at the center than that of any other quadrant. Compared to untreated cells, hyper-acetylation or ATP depletion did not substantially change this pattern, thus confirming previous results (Fig. 1) that central/peripheral heterogeneities in chromatin fluctuations are robust to metabolic changes and acetylation of histones.

### H4 histones interact with dBD except in cells both hyper-acetylated and ATP-depleted

To better understand the effect of chromatin condensation and its fluctuations on accessibility of protein factors, we sought to assess the interaction of the TAFII250-double bromodomain (dBD) with acetylated H4 histones. First, to ensure that we were not in limiting concentrations of dBD, we quantified the presence of excess dBD by FCS (Fluorescence fluctuation correlation spectroscopy) on dBD-GFP. In cells exogenously expressing dBD-GFP, FCS showed the protein sorted in two populations: about 12% of dBD-GFP exhibited an apparent diffusion coefficient of 0.3 μm^2^s^−1^, consistent with that of a population interacting with an immobile ligand, the rest exhibited an apparent coefficient of about 4 μm^2^s^−1^, consistent with a protein diffusing much more freely. Thus, dBD-GFP is present in large excess in those conditions (Table S1). Since TAFII250 is known to interact with acetylated lysines of H4 histones, we sought to also ensure that hyper-acetylation would not deplete the excess of dBD-GFP. In this condition, the slow population exhibited an apparent diffusion coefficient indistinguishable from that of the slow population in untreated condition and its fraction increased to about 20% of total dBD-GFP. This is consistent with an increase in acetylated H4 histones capturing a larger amount of dBD-GFP, but not to the point of free dBD depletion.

We thus proceeded to directly assess the interaction between EGFP-H4 and dBD-mCherry. Compared to cells expressing EGFP-H4 alone, cells expressing both EGFP-H4 and dBD-mCherry exhibited a nuclear map of EGFP-H4 *τ*̃ shifted to lower values (Fig. 3A,B). Significantly, both peripheral and central regions of the nucleus exhibited *τ*̃ lower in the presence of dBD-mCherry than in its absence (Δ*τ*̃ > 0.04ns and 0.02ns, respectively), indicating that FRET, and thereby an interaction occurs between EGFP-H4 and dBD-mCherry, and apparently to a larger extent at the periphery (Fig. 3C). Similarly in hyper-acetylation condition, both peripheral and central regions of the nucleus exhibited *τ*̃ lower in the presence of dBD-mCherry (Δ*τ*̃ > 0.06ns and 0.04ns, respectively). In ATP depleted cells, *τ*̃ also exhibited a drop in the presence of dBD-mCherry both at the periphery and the center (Δ*τ*̃ > 0.05ns and 0.03ns, respectively). This shows that accessibility of dBD to H4 histones does not require ATP, just as chromatin fluctuations do, and still occurs in condensed chromatin condition (see Fig. 1). However, in hyper-acetylated and ATP-depleted cells, the drop in *τ*̃ was dramatically reduced of near an order of magnitude both at the periphery and the center (Δ*τ*̃ < 0.01ns and 0.001ns, respectively) such that *τ*̃ distributions at the center with or without dBD were indistinguishable, in the only condition that also led to a substantial decrease in chromatin fluctuations (see above). Finally, the drops in *τ*̃ were in all conditions larger at the periphery than in the center of the nucleus, just as fluctuations were (Fig. 1).

**Figure 3:**
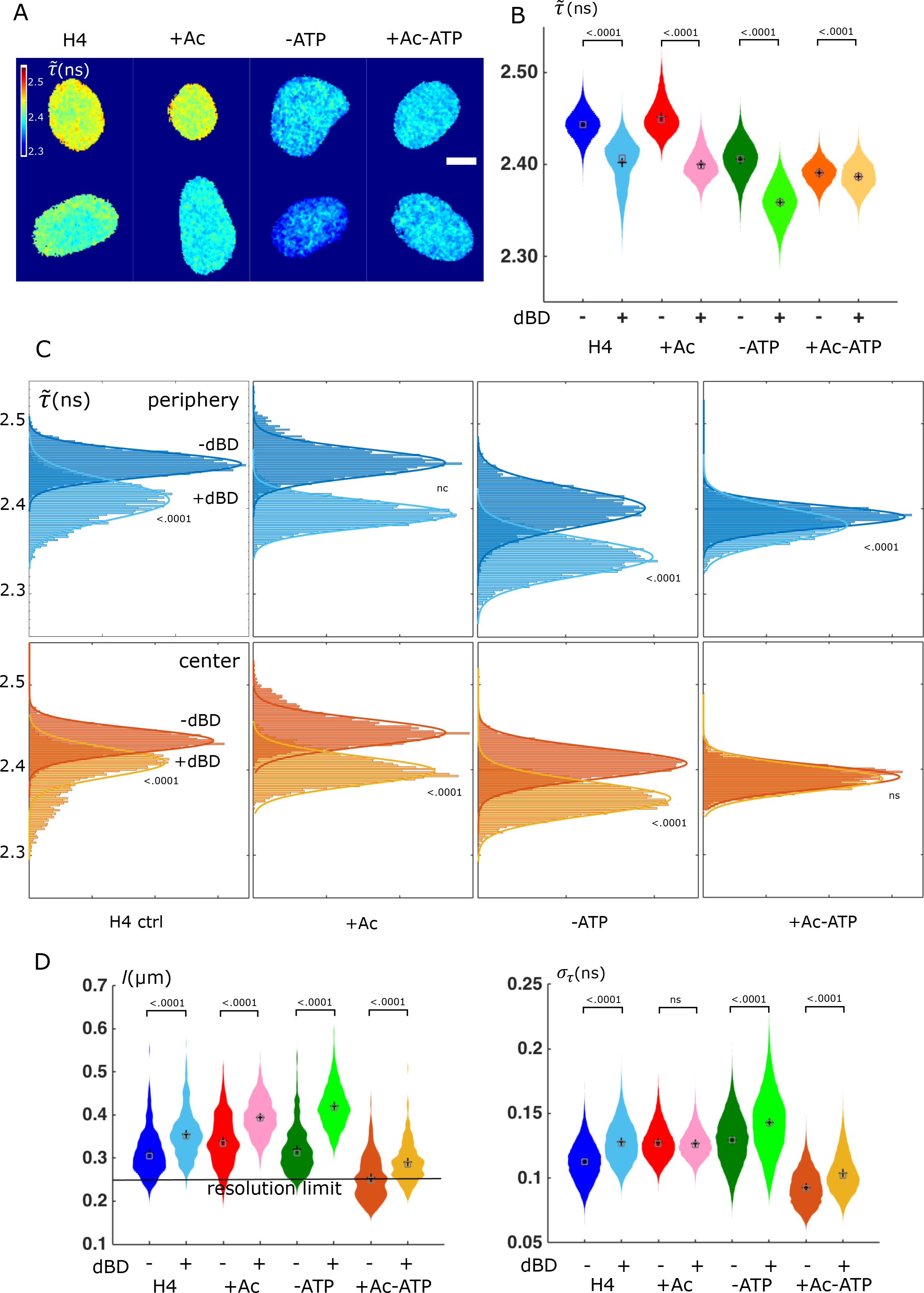
**A:** Typical images of time-averaged fluorescence lifetime *τ*̃ in HEK cells expressing EGFP-H4 in untreated (H4), hyper-acetylation (+Ac), ATP-depletion (- ATP), or both (+Ac-ATP) conditions, in the absence (top) or presence (bottom) of dBD-mCherry. Bar = 5 μm. **B:** Time-averaged fluorescence lifetime *τ*̃ distributions in conditions above. Same data as in Fig. 1 for absent dBD-mCherry, otherwise EGFP-H4 (16 474 pixels, 8 cells); +Ac (19 146 pixels, 9 cells); -ATP (24 432 pixels, 13 cells); +Ac-ATP (21 026 pixels, 8 cells). Black cross: mean, green box: median. Kruskal-Wallis test. **C:** Time-averaged fluorescence lifetime *τ*̃ distributions in conditions above for peripheral and central pixels and their Gaussian fits. Same as in Fig. 1 for absent dBD m-Cherry. Extra sum-of-squares F test with same fit for both distributions as null hypothesis. ns=not significant, nc= not convergent (for a same fit for both distributions). **D:** Correlation length *l* distributions in conditions above. Same data as in Fig. S1-D for absent dBD-mCherry, otherwise EGFP-H4 (16 474 pixels, 8 cells); +Ac (19 146 pixels, 9 cells); -ATP (24 432 pixels, 13 cells); +Ac-ATP (21 026 pixels, 8 cells). Black cross: mean, green box: median. Kruskal-Wallis test. **E:** Distributions of fluorescence lifetime standard deviation *σ*_*τ*_ in conditions above. Same data as in Fig. 1 for absent dBD-mCherry, otherwise EGFP-H4 (16 474 pixels, 8 cells); +Ac (19 146 pixels, 9 cells); -ATP (24 432 pixels, 13 cells); +Ac-ATP (21 026 pixels, 8 cells). Black cross: mean, green box: median. Kruskal-Wallis test. ns= not significant.

### Chromatin accessibility domains are larger than condensation domains and accessibility fluctuations are restricted by hyper-acetylation in an ATP-dependent fashion

In the presence of dBD, similarly to Δ*τ*̃ reporting chromatin accessibility, *l* and Δ*σ*_*τ*_ may report the typical size of accessible domains and the occurrence of temporal fluctuations of accessibility, respectively.

In the presence of dBD, *l* was higher than in its absence in all conditions examined (Δ*l* >36nm), which indicates that accessible chromatin domains are larger than condensation domains (Fig. 3D). The spontaneous fluctuation condition (-ATP) exhibited the largest accessible domains (*l*=420nm) and the decrease in accessible domain size was smaller upon addition of ATP and hyper-acetylation together (Δ*l*=25nm) than upon any of the two (Δ*l*=64nm and Δ*l*=130nm, respectively). Moreover, the discrepancy between condensation and accessible domains sizes also decreased less upon addition of ATP and hyper-acetylation (Δ*l*=36nm) than upon any of the two (Δ*l*=43nm and Δ*l*=56nm, respectively). Altogether, this indicates the existence of hyper-acetylation and ATP-dependent mechanisms partly independent of chromatin condensation domains size that individually restrict accessible chromatin domains size but are out-competed by ATP- and hyper-acetylation-dependent mechanisms.

In the presence of dBD, *σ*_*τ*_ was substantially higher than in its absence in all (Δ*σ*_*τ*_ > 0.011ns) but the hyper-acetylated condition (Fig. 3E). This indicates that significant accessibility fluctuations occur in addition to condensation fluctuations. Nevertheless, the lack of significant difference in *σ*_*τ*_ with and without dBD in hyper-acetylated cells (+Ac) reveals that in this condition, accessibility fluctuations are strictly regulated such that overall *σ*_*τ*_ amplitudes do not exceed that of condensation fluctuations.

### Chromatin condensation fluctuations predict chromatin accessibility

So far, our results support that chromatin accessibility is granted provided sufficient chromatin fluctuations occur, regardless of chromatin steady-state condensation. To characterize this more quantitatively, we sought to assess the correlation between *σ*_*τ*_ or *τ*̃ on the one hand, and the extent of FRET on the other end, throughout all experimental conditions. We found that the mean FRET transfer rate *k*FRET (see SI Material and Methods) significantly correlated with *σ*_*τ*_ (r²=0.8783, p<0.001) while it did not with *τ*̃ (r²=0.421, p>0.08) (Fig. 4A). These results support that chromatin accessibility is sensitive to chromatin condensation fluctuations amplitudes rather than chromatin condensation level.

**Figure 4:**
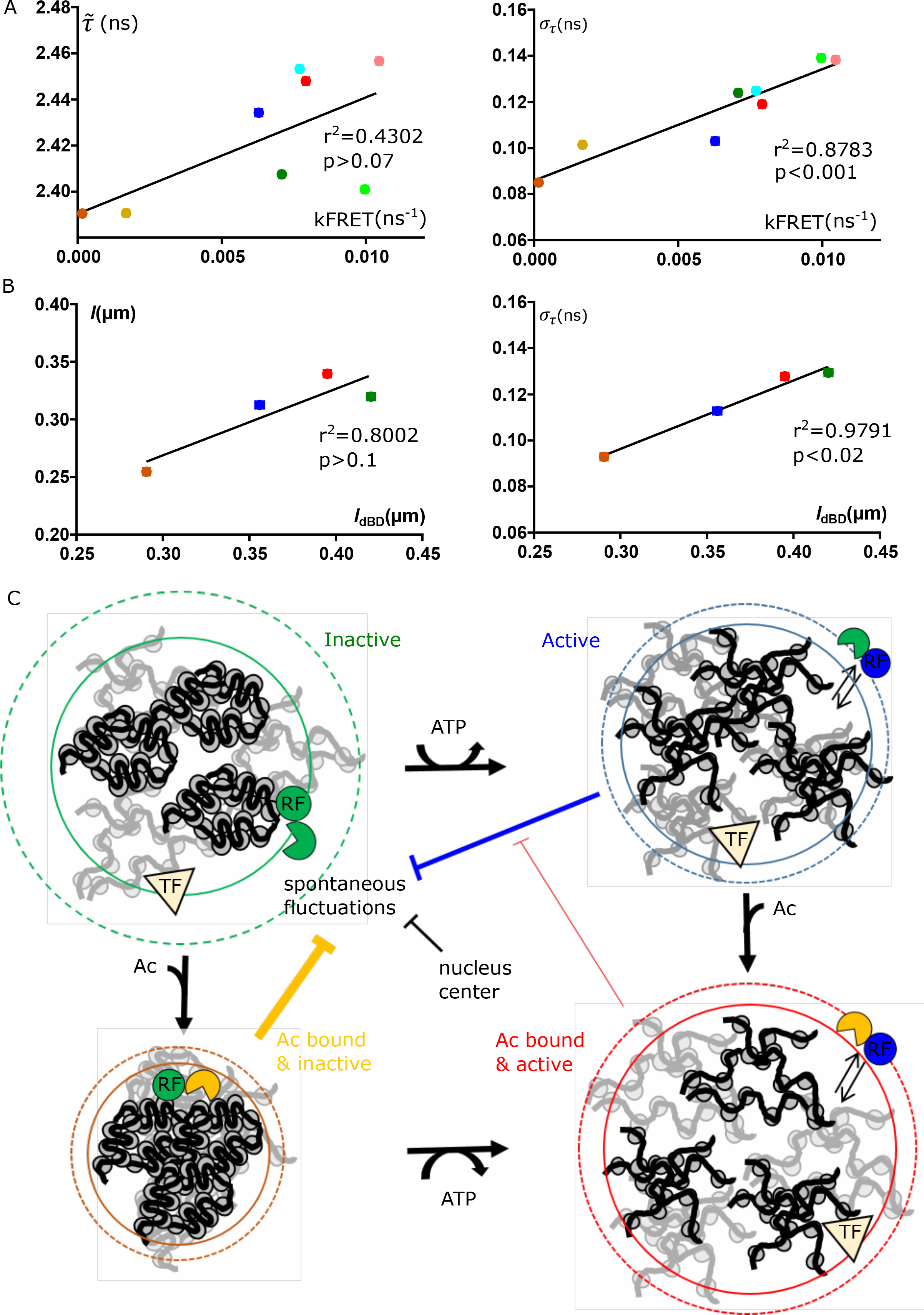
**A:** Time-averaged fluorescence lifetime *τ*̃ (left) and standard deviation *σ*_*τ*_ (right) as a function of *k*FRET in all conditions (blue: H4, red: +Ac, green: -ATP, brown: +Ac-ATP) and sub-nulcear regions (dark: center, pale: periphery). Data are mean +/- SEM of Gaussian fits from data shown in Figs. 1C and F. Solid lines are linear fits, correlation coefficients displayed. Extra sumof-squares F test with slope=0 as null hypothesis. **B:** Fluorescence lifetime spatial correlation length *l* (left) and standard deviation *σ*_*τ*_ (right) as a function of spatial correlation in the presence of dBD *l*_dBD_ in all conditions (blue: H4, red: +Ac, green: -ATP, brown: +Ac-ATP). Data are mean +/- SEM of whole nucleus distributions from Figs. 1 and SX). Solid lines are linear fits, correlation coefficients displayed. Extra sum-of-squares F test with slope=0 as null hypothesis. **C:** Mechanisms regulating spontaneous fluctuations of chromatin condensation. TF: transcription factor, RF: ATP-dependent and acetylation-binding remodeling factor and its state (active/inactive, Ac bound or not). Plain circle: same-condensation domain size, dashed circle: same accessibility domain size. See text for details.

To substantiate this further, we assessed the correlation of accessible domain size *l*_dBD_ with condensation domain size *l* and with condensation fluctuation amplitudes *σ*_*τ*_ (Fig. 4B). Consistently, we found that accessible domain size significantly correlated with condensation fluctuation amplitudes (r²=0.971, p<0.02) whereas it did not with condensation domain size (r²=0.8002, p>0.1).

Altogether, these results show that chromatin accessibility is predicted by chromatin condensation fluctuations rather than steady-state.

## DISCUSSION

### Fluorescence lifetime of GFP tagged on histones reports chromatin condensation

The fluorescence lifetime of a fluorophore depends on its local environment, which may affect both radiative and non-radiative decays. While the fluorescence lifetime of GFP was shown to be insensitive to viscosity or pH (25, 26), unlike that of DNA binding dyes (27), its dependency on the refractive index *n* (28) was quantitatively verified in solution (29) and previously used with tagged proteins to probe *n* in live cells (30, 31).

Applying the published calibration curve (29) to our results, *τ*̃ gives for EGFP-H4 in untreated condition *n*=1.398 ± 0.002 and 1.405 ± 0.003 for nuclear periphery and center, respectively (from Fig. 1, see SI Materials and Methods). These values are in remarkable agreement with mean nuclear refractive index measurements (32). Of note, the fluorescence lifetime sensitivity to *n* is estimated to exponentially drop by one half about 1µm away from the fluorophore (33), such that the correlation length of *τ* we observe (Fig. S3) is in the reach of the fluorescence lifetime sensitivity to local refractive index changes. Thus, spatial heterogeneities of *τ* may hide heterogeneities in chromatin condensation even smaller and sharper than we could see, which would comprise a few thousand nucleosomes.

In summary, the spatial and temporal variations in fluorescence lifetime EGFP-H4 are overall consistent with spatial and temporal change of refractive index corresponding to chromatin condensation changes.

### Relationship with chromatin condensation and its fluctuations assessed by other methods

Accumulating data on chromatin reporters demonstrate that chromatin in the interphase nucleus undergoes various types of fluctuations. Localization microscopy provides chromatin density fluctuations at the nm scale (34, 35). Single particle tracking shows chromatin motions over submicron distances as fast as within the 10 msec timescale (15–17). At the opposite end, displacement correlation spectroscopy shows motions at the 1-10 sec timescales over several microns (18).

Here, fluorescence lifetime fluctuations are in an intermediate range (sec timescale and 100nm length scale) (Figs. 1, S1), thus report intermediate level of chromatin fluctuations. Of note, the spatial correlation length we measure corresponds to a volume that would contain about 4.10^4^ nucleosomes assuming a nucleosome concentration of 100 μM (36).

### Distinct mechanisms regulate chromatin condensation and its fluctuations

We have evidenced three remarkable features of chromatin condensation fluctuations. First, these fluctuations are spontaneous, since ATP depletion results in an increase in fluctuation amplitude (Fig. 1). In agreement, in vitro reconstitution studies have shown that individual, and arrays of nucleosomes undergo spontaneous conformational fluctuations across the µsec to min timescales (37–39). Moreover, fixed cells also exhibit nucleosome fluctuations (17). We propose that the activity of ATP-dependent chromatin remodelers is to some extent detrimental to spontaneous chromatin condensation fluctuations (Fig. 4C). This is stark contrast with the regulation of chromatin condensation, since chromatin de-condensation is ATP-dependent (Fig. 1). Consistently with this, ATP-depletion promotes fluctuations in condensed regions in particular (see *σ*_*τ*_ - *τ*̃ anti-correlation in Fig. 2). Thus chromatin de-condensation and its fluctuations are supported by fundamentally distinct mechanisms.

Second, chromatin fluctuations are intrinsically dependent on chromatin sub-nuclear localization, since their amplitude is consistently higher at the nuclear periphery regardless of acetylation or metabolic conditions (Fig. 1). This shows that radial heterogeneities in chromatin fluctuations cannot be explained solely from radial heterogeneities in histone acetylation and ATP-dependent remodeling activity. Thus, sub-nuclear spatial cues constitute a distinct pathway regulating chromatin fluctuations. This is again in stark contrast with chromatin condensation, which although exhibits radial heterogeneity too, requires ATP to do so (Fig. 1).

Third, we identify (1) acetylation-independent, ATP-dependent mechanisms and (2) acetylation-dependent, ATP-independent mechanisms that each decrease spontaneous fluctuations, but these are out-competed by (3) acetylation- and ATP-dependent mechanisms restoring fluctuations closer to their spontaneous level (Fig. 1). We propose that (1) ATP hinders spontaneous fluctuations through activation of acetylation-independent, transcriptional repressor-associated remodeling complexes, (2) factors binding to acetylated chromatin restrict spontaneous chromatin condensation fluctuations in the absence of ATP, but that (3) ATP promotes the release of these factors, thereby relieving part of the hindrance (Fig. 5C). Indeed, ligands of acetylated chromatin are widespread among ATP-dependent remodeling complexes (24), and the release from chromatin of one of them was previously found to require ATP (40). Furthermore, in hyper-acetylated condition, the presence of ATP may in addition facilitate acetylation propagation on adjacent nucleosomes by acetyltransferases (41), and thereby fluctuations. In line with this, hyper-acetylation promotes chromatin de-condensation and increases the size of chromatin same-condensation domains provided ATP is present, and the opposite, when ATP is absent (Fig. S1). Mechanisms of interplay between chromatin acetylation and ATP-dependent remodeling are currently being revealed (42, 43). Future studies will address the identification of the factors involved here.

### Mechanisms underlying chromatin accessibility

Genome accessibility has mostly been assessed with biochemical approaches on isolated nuclei on the basis of DNA sensitivity to some enzymatic activity (digestion, methylation…). (44). Besides, a number of fluorescence imaging modalities has been applied to assess intranuclear milieu permeability to inert fluorescent particles or chromatin interactants (11–14, 17, 45). Of note, some of these studies have showed that chromatin is almost as permeable regardless of its condensation state.

Beyond mere intranuclear permeability or the mobility of chromatin interactants, we probe here by FRET the direct interaction of chromatin with a domain of transcription factors. Our approach thus provides a direct and quantitative assessment of chromatin accessibility to biologically relevant molecules. Our results show that conditions that elicit chromatin condensation do not suffice to impair chromatin accessibility (Fig. 3). Consistently, chromatin condensation domains do not restrict in size chromatin accessibility domains (Fig. 3). This extend previous observations of condensation-independent chromatin permeability. We show instead that chromatin condensation fluctuation level predicts accessibility level and domains size better than condensation level and domain size do (Fig. 4). That condensed chromatin may exist in distinct fluctuation states with distinct accessibilities to transcription factors may explain inconsistencies between previously reported condensed chromatin permeability.

The mechanisms underlying chromatin accessibility to acetylated histones are directly related to those affecting condensation fluctuations. Notably, accessibility is spontaneous, as it occurs in absence of ATP, and it is distinct from acetylation level, since hyper-acetylation severely impairs accessibility in the absence of ATP (Fig. 1). Thus, acetylation level is alone a poor indicator of accessibility. Consistently, in the presence of ATP, the accessibility increase upon hyper-acetylation is somewhat modest (Fig. 1) when compared to the extent of hyper-acetylation (Fig. S1). This supports that, even with ATP, only a fraction of acetylated histones is accessible within accessible domains. We propose that while ATP allows the release of acetylation- and ATP-dependent binding factors, thereby condensation fluctuations, and subsequently accessibility to a fraction of acetylated histones, another fraction remains inaccessible due to other binding factors that prevent condensation fluctuations. Whether this fraction organizes as in previously reported sub-diffraction domains remains to be determined (19). Other ATP- and hyper-acetylation-dependent mechanisms may also account for the limited accessibility of acetylated histones, as fluctuations of accessibility are restricted to a level indistinguishable from condensation fluctuations in the presence of both ATP and hyper-acetylation.

In conclusion, we show that the critical determinant for chromatin accessibility is its fluctuations, rather than its condensation. Moreover, these fluctuations are spontaneous, intrinsically dependent on sub-nuclear localization, and under the control distinct and competing mechanisms dependent on acetylation, ATP, and both.

## MATERIALS AND METHODS

HEK 293 cells transiently expressing fluorescently tagged proteins were monitored on a two-photon FLIM system 2 days after transfection and after 24hrs or 2hrs after drug treatment, depending on experiment. Image and data analyses were performed with MATLAB (MathWorks) and Prism (GraphPad). Full materials and methods are available in SI Materials and Methods.

## ACKNOWLEDGEMENTS

We thank Sophie Polo for critical reading of the manuscript. We thank Mathieu Coppey for custom-built MATLAB routines, discussions and critical reading of the manuscript. This work was partly supported by grants from IBiSA and Fondation Recherche Médicale, and by the Centre National de la Recherche Scientifique. We acknowledge the ImagoSeine facility of the Institut Jacques Monod, member of France BioImaging infrastructure supported by ANR-10-INBS-04.

## SI MATERIALS AND METHODS

### Plasmid constructs

Fluorescent fusion proteins were cloned in pEGFP-C1 (Clontech, Mountain View, CA) and pmCherry-C1. To generate pmCherry-C1, the mCherry coding sequence was transferred from pRSETB-Cherry (a generous gift from Dr. Tsien, University of California at San Diego) into a Clontech vector backbone. H4 cDNA (IMAGE: 2130477, ResGen) was cloned in pEGFP-C1 vector using Xhol and SalI sites. The dBD sequence (containing the amino acids 1207–1872 of TAFII250) was obtained from Kanno et al. (1) and cloned in the pmCherry-C1 or in the pEGFP-C1 vector using EcoRI and KpnI sites.

### Cell culture and treatments

HEK 293 cells were cultured in Dulbecco’s modified Eagle’s medium containing 10% fetal bovine serum (PAA Laboratories GmbH, Pasching, Austria). The cultures were incubated at 37°C in a humidified atmosphere of 5% CO_2_. HEK293 cells were seeded on 32-mm round glass coverslip at a density of 2×10^5^ cells. At ~70% of confluence, cells were transfected with a total amount of 1 μg of expression vectors using Nanofectin I (PAA Laboratories, Inc). Twenty-four hours later, coverslips were mounted in an open observation chamber with DMEM-F12 without Phenol red, B12 vitamin and Riboflavin, and supplemented with 20 mM HEPES and L-Glutamine (PAA). Sodium butyrate (NaB), sodium azide (Az), and deoxyglucose (DG) were obtained from Sigma Aldrich (Stenheim, Germany). For NaB treatment, cells were incubated 24 hours after transfection with 2.5 mM of NaB for 24 hours before imaging. For ATP depletion, cells were incubated with 10 mM Az and 10 mM DG for 2 hours before imaging. Histone acetylation was verified after Acid-Urea-Triton Polyacrylamide gel electrophoresis (15% acrylamide, 0.1% M-bisacrylamide, 6 M urea, 5% acetic acid, 8 mM triton X-100, 0.5% TEMED, 0.00025% riboflavin) by western blot with an anti-H4 primary antibody (07-108, Upstate, France), and peroxidase-conjugated secondary antibody (Amersham ECL Western blotting kit) (Fig. S1).

### Fluorescence lifetime acquisition

Cells were imaged on a two-photon FLIM system (TriM-FLIM) that combines multifocal multiphoton excitation (TriMscope) and a fast-gated CCD camera (PicoStar), all controlled by the IMspector software (LaVisionBiotec, Bielefeld, Germany), as described elsewhere (2). Briefly, two-photon multifocal excitation is carried out using the TriMScope connected to an inverted microscope (IX 71, Olympus, Tokyo, Japan). A mode-locked Ti:Sa laser tuned at 950 nm for the excitation of enhanced green fluorescent protein (EGFP) (MaiTai Spectra Physics, Evry, France) is split into 64 beams with a 50/50 beam splitter and mirrors. The beams pass through a 2000 Hz scanner before illuminating the back aperture of an infrared water immersion objective (60X/NA 1.2, Olympus, Tokyo, Japan). A line of foci at the focal plane is scanned across the sample, generating a pseudo wide-field illumination. The x-y spatial resolution, measured by point spread function, is r = 250 nm. The spatial resolution along the optical axis is given by the diffraction of 960 nm laser beam in z direction focused through the NA 1.2 objective, i.e. 1400 nm. A wheel of spectral filters (520DF40, Chroma Technology, for EGFP) is used to select the fluorescence imaged onto a fast-gated light intensifier connected to a CCD camera (PicoStar). The gate of the intensifier is set to 2 ns. The intensifier acts as a very fast shutter for each delay from 0 to 10 ns along the fluorescence decay. 5 gates are necessary and sufficient to calculate the mean, instantaneous fluorescence lifetime (*τ*)of EGFP (2), by applying: *τ* = ΣΔ*t*_i_. *I*_i_/Σ*I*_i_, where Δ*t*_i_ corresponds to the time delay after the laser pulse of the i^th^ gated image acquired and *I*_i_ to the pixel intensity map in the i^th^ gated image. The acquisition time of the CCD camera was set to 3 s, which determines the time resolution of a *τ* image. The time-average of *τ* (*τ*̃ in the main text) and its temporal standard deviation (*σ*_*τ*_) were calculated from 100 consecutive images for each pixel using ImageJ (W. S. Rasband, ImageJ, U.SNational Institutes of Health, Bethesda, Maryland, http://rsb.info.nih.gov/ij/) (Fig. S2). The steady-state fluorescence intensity is the sum of the five gated images intensities (ΣI_i_) averaged over 4 consecutive images.

### Fluorescence lifetime analysis

#### Data management and analysis

Nuclear pixels were selected based on EGFP-H4 steady-state fluorescence intensity thresholding. Nuclear pixel coordinates were used to calculate the smallest distance of each nuclear pixel to the nucleus border (*d*), the mean distance per nucleus of a nuclear pixel to the border (*d*̃) and its standard deviation (*σ_d_*), to retrieve the deviation of each nuclear pixel from the mean distance to the border (*δ* = (*d* − *d*̃)/ *σ*_d_, which typically ranged from −1.5 (close to the border) to 3 (far from the border). Peripheral pixels were defined as *δ* ≤ −1 and central pixels as *δ* ≥ +1 (Fig. S4A). Fluorescence lifetime data (*τ*, *τ*̃, *σ*_*τ*_) of every nuclear pixel in 7-15 cells were collected from at least 3 independent experiments for each condition and their distributions displayed in ‘violin’ plots, histograms according to *σ*_*τ*_, or 2D-plots depending on figures. All of the above was performed in MATLAB (MathWorks).

#### Refractive index determination

The refractive index *n* is related to the fluorescence lifetime *τ* by the equation (3):

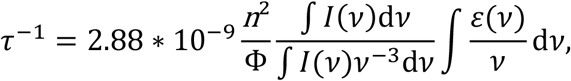

in which Φ is the quantum yield, *I* is the fluorescence emission intensity at the wave number *ν* and *ε* is the extinction coefficient. The mean values of *τ* obtained from the Gaussian fit of the distributions at nuclear periphery and center were reported on the calibration curve published elsewhere (4), to obtained the refractive indices *n* at nuclear periphery and center, respectively.

#### Spatial correlation analysis

The instantaneous fluorescence lifetime fluctuation spatial autocorrelation function of the FLIM image recorded at time *t* in a time series is given by:

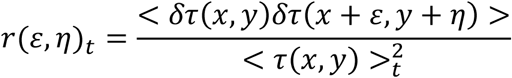

where the angular brackets denote spatial averaging over the image, ε and η are spatial lag variables corresponding to pixel shifts of the image relative to itself in the *x* and *y* directions and *δ*τ** the difference between **τ** and its spatial mean at *t*. Images are analysed by a MATLAB routine (a generous gift from David L. Kolin and Paul W. Wiseman) and the correlation functions calculated using Fourier methods and then fit to a 2D Gaussian, which half width at half maximum is the correlation length *l*.

#### FRET quantification

The mean transfer rate is defined by *k*FRET = 1/*τ*_D_ + 1/*τ*_DA_, in which *τ*_D_ and *τ*_DA_ are the donor fluorescence lifetime in the absence and in the presence of acceptor, respectively (5). For each experimental condition (H4, +Ac, -ATP, +Ac-ATP, for nucleus periphery and center), *τ*_D_ and *τ*_DA_ were obtained from Gaussian fits of *τ*̃ distributions in the absence and in the presence of dBD, respectively.

#### Statistical Analysis

Comparisons between treatments were performed with (unpaired, two-tailed) Kruskal-Wallis tests. Comparisons between peripheral and central distributions were performed on Gaussian fits with extra sum-of-squares F tests and the null hypothesis that both distributions are better fitted together by a single Gaussian rather than one each. Correlations were tested with extra sum-of-squares F tests and the null hypothesis that slopes of linear fits were 0. All of the above was performed in Prism V (GraphPad).

### FCS measurements

FCS measurements were carried out on a confocal time-resolved microscope MicroTime 200 (Picoquant, Berlin, Germany) as described elsewhere (6). dBD-EGFP was excited at 470 nm by a pulsed diode laser (Picoquant) through an Olympus UPLSAPO water objective (60X/NA 1.2), for a 0.4 fL excitation volume. Fluorescence emission passes through a dichroic mirror (DM 470 nm, Picoquant) and is focused on a pinhole in front of an avalanche photodiode (SPADs 14, Perkin Elmer) through the emission filter FF01-525/50 (Picoquant). The TimeHarp 300 PC board (PicoQuant) acquires during 60 sec single photon-counting time traces analyzed with the Symphotime software (PicoQuant). A two-species Brownian diffusion model was used to fit the fluorescence fluctuation auto-correlation function and retrieve amplitudes and apparent diffusion coefficients.

### FRAP experiments

FRAP (Fluorescence Recovery After Photobleaching) experiments were carried out on a Leica DMI6000 microscope equipped with a Yokogawa CSU22 spinning-disk head, a Leica Plan APO 63X/1.2 NA water objective and an EMCCD camera (Photometrics QuantEM). A FRAP Head (Roper scientific) equipped with a 473 nm diode laser (100 mW) was used to perform point bleaching. Pre-bleach and post-bleach image acquisitions were performed with a 491 nm diode laser (50 mW). Fixed cells expressing the same EGFP constructs were used to determine the 473nm laser power required to bleach all the fluorescence, and the 491nm laser power required not to bleach fluorescence through imaging.

---

**Figure S1:**
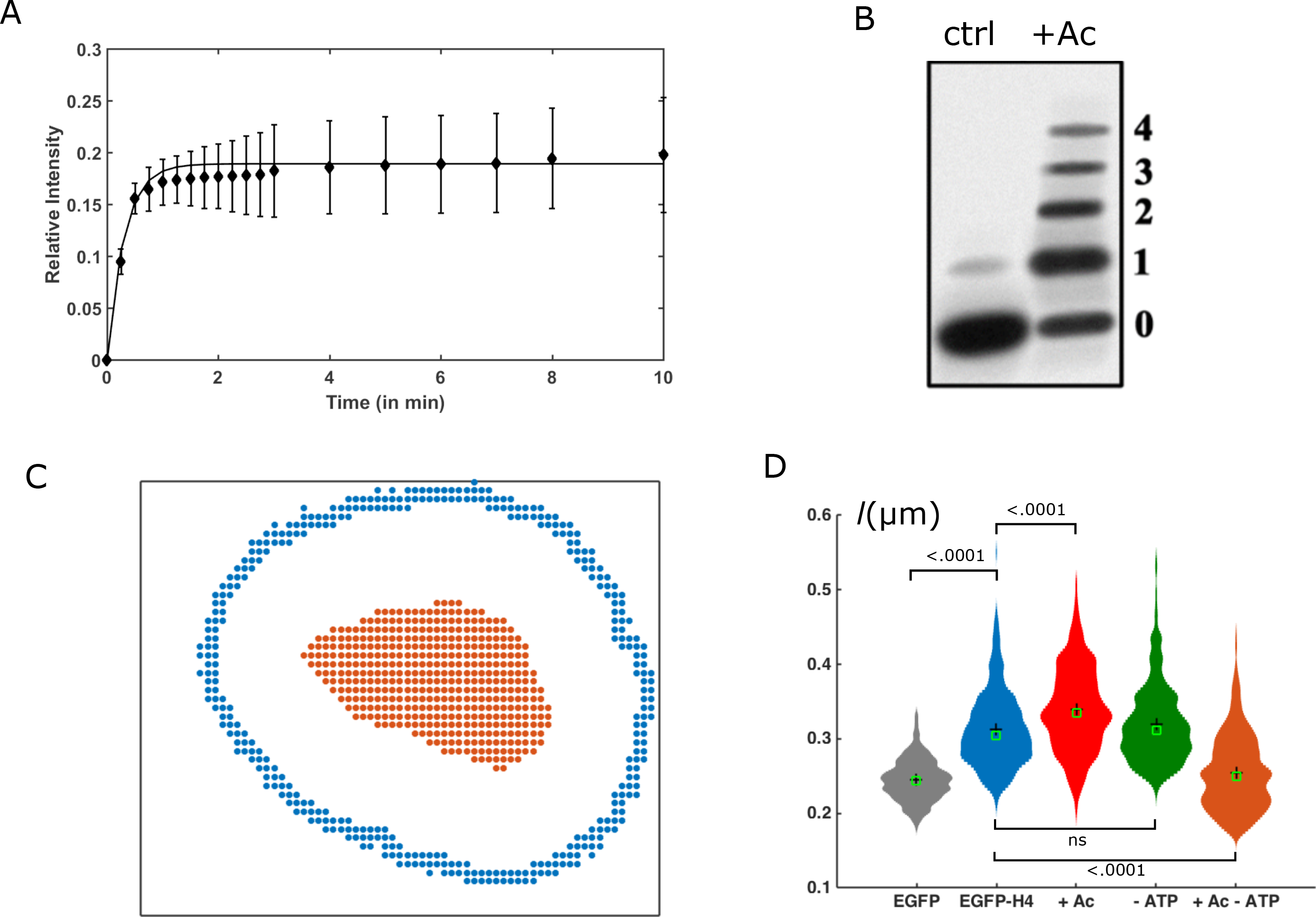
**A:** Fluorescence recovery after photobleaching of EGFP-H4 (mean +/- SEM, n=30 cells. **B:** Western blot of histones in untreated and hyper-acetylation conditions. Hyper-acetylation results in the enrichment of mono-, di-, tri- and tetra-acetylated histone fractions from 7 to 35%, and 0 to 19, to 11, and to 8%, respectively. **C:** Typical segmentation of peripheral (blue *δ* ≤ −1) and central (*δ* ≥ 1) nuclear pixels. See SI Materials & Methods. **D:** Distributions of spatial correlation length *l* of *τ* in EGFP (100 images, 7 cells), control (H4 ctrl, 100 images, 13 cells), hyper-acetylation (+Ac, 100 images, 8 cells), ATP-depletion (- ATP, 100 images, 13 cells), or both (+Ac-ATP, 100 images, 8 cells) conditions. Black cross: mean, green box: median. Kruskal-Wallis test. ns=not significant.

**Figure S2:**
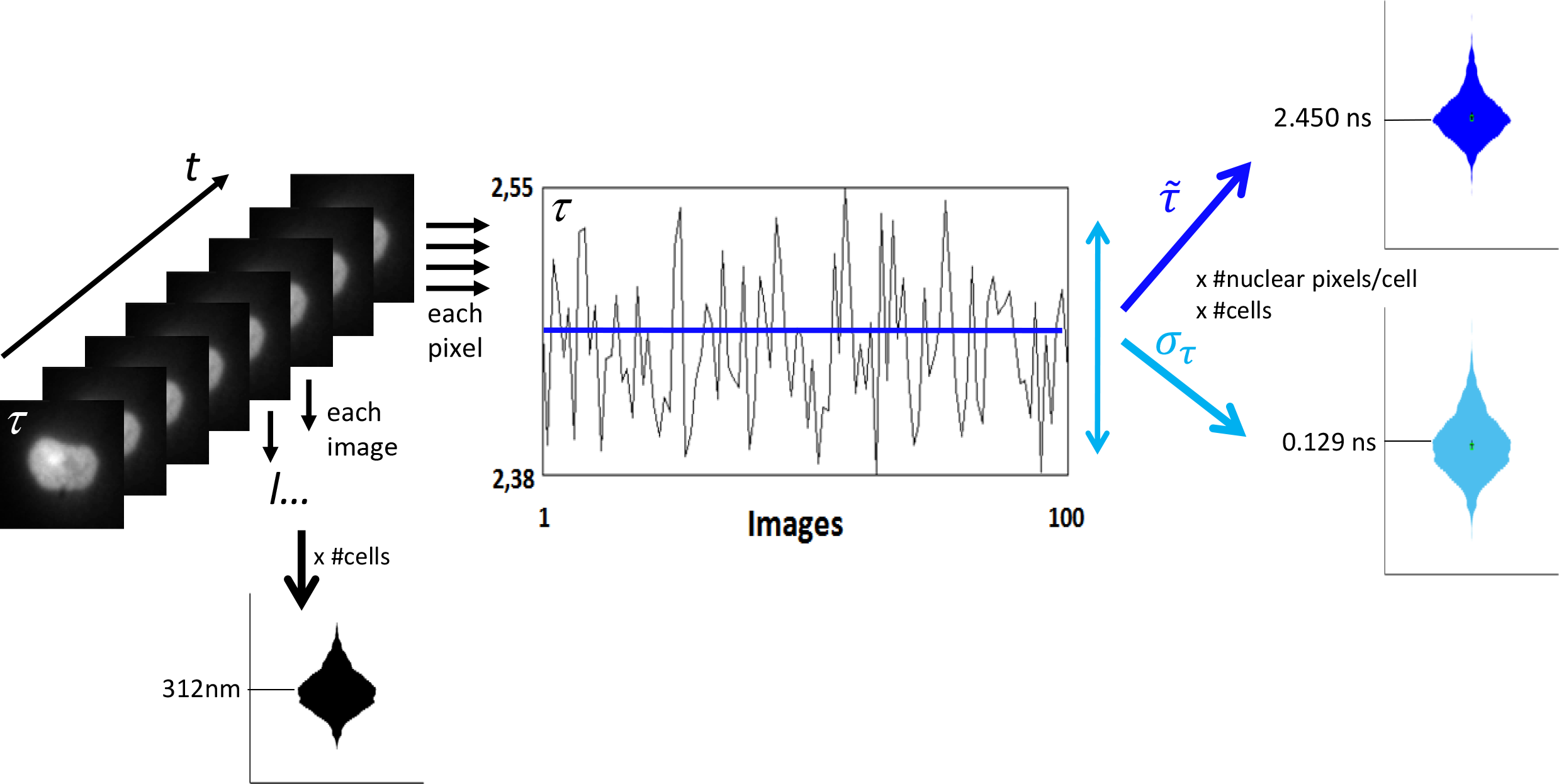
Data acquisition workflow. For each cell in each condition, 100 consecutive images of instantaneous fluorescence lifetime *τ* are acquired (3 sec time interval). From each image is retrieved the spatial correlation length *l*, and each pixel (x, y coordinates) across the 100 images the time-averaged fluorescence lifetime *τ*̃ and the standard deviation *σ*_*τ*_.

**Table S1:** Apparent diffusion coefficients and proportions of EGFP and EGFP-dBD mobile species in nuclei (mean+/-SD).

| | Diffusion Coefficient ($\mu\text{m}^2/\text{s}$) | Proportion (%) |
| --- | --- | --- |
| EGFP | 10.04+/-1.73 | 100 |
| EGFP-dBD | 0.28+/-0.31 | 12.3+/-6.7 |
|  | 3.84+/-0.8 | 87.7+/-6.7 |
| EGFP-dBD (+Ac) | 0.32+/-0.16 | 21.4+/-12.2 |
|  | 4.23+/-1.26 | 78.6+/-12.2 |

Author contributions
N.A., S.P.-P., M.T. performed experiments; N.A., S.P.-P., M.T. Developed methodology; N.A. N.B. and M.C.-M. analyzed data; M.C.-M. supervised research; N.B. and M.C.-M. wrote the manuscript.

